# Maternal alum-adjuvanted recombinant HIV Env vaccine does not enhance autologous virus neutralization in HIV-infected pregnant women

**DOI:** 10.1101/2019.12.11.873018

**Authors:** Eliza D. Hompe, Jesse F. Mangold, Joshua A. Eudailey, Elena E. Giorgi, Amit Kumar, Erin McGuire, Barton F. Haynes, M. Anthony Moody, Peter F. Wright, Genevieve G. Fouda, Feng Gao, Sallie R. Permar

**Affiliations:** Duke Human Vaccine Institute, Duke University Medical Center, Durham, NC, USA; Los Alamos National Laboratory, Los Alamos, NM, USA; Department of Medicine, Duke University Medical Center, Durham, NC, USA; Department of Pediatrics, Duke University Medical Center, Durham, NC, USA; Department of Pediatrics, Dartmouth-Hitchcock Medical Center, Lebanon, NH, USA

**Keywords:** Human immunodeficiency virus, mother-to-child transmission, maternal vaccination, HIV envelope, autologous virus neutralization

## Abstract

Preventive strategies beyond ART will be required to end the pediatric HIV epidemic. A maternal vaccine capable of boosting neutralizing antibody (nAb) responses against circulating viruses in HIV-infected pregnant women could effectively decrease mother-to-child transmission of HIV. However, it is not known if an HIV envelope (Env) vaccine administered to infected pregnant women can enhance autologous virus neutralization.

Here, we assessed autologous virus nAb responses in maternal plasma samples obtained from AIDS Vaccine Evaluation Group (AVEG) Protocols 104 and 102, historical Phase I safety and immunogenicity trials of recombinant HIV Env subunit vaccines in HIV-infected pregnant women (NCT00001041). AVEG 104 participants were randomized to receive 300 µg Env subunit MN recombinant gp120 with alum adjuvant or alum alone. AVEG 102 participants were randomized to receive 640 µg Env subunit recombinant gp160 or placebo. HIV Env-specific maternal plasma binding and neutralizing responses were characterized before and after vaccination in 15 AVEG 104 (n=10 vaccinee, n=5 placebo) and 2 AVEG 102 (n=1 vaccinee, n=1 placebo) participants. Single genome amplification (SGA) was used to obtain HIV *env* gene sequences from autologous viruses for pseudovirus production in pre- and post-vaccination plasma of HIV-infected pregnant vaccinees (n=6 gp120, n=1 gp160) and placebo recipients (n=3).

We detected an increase in MN gp120-specific IgG binding in the vaccinee group between the first immunization visit and the last visit at delivery (p=0.027, 2-sided Wilcoxon test). However, no difference was observed in the neutralization potency of maternal plasma collected at delivery against autologous viruses isolated from early or late pregnancy. Thus, maternal vaccination with gp120/160 did not boost maternal autologous virus nAb responses. Immunization strategies capable of more potent B cell stimulation will likely be required to effectively boost autologous virus nAb responses in pregnant women and synergize with ART to further reduce infant HIV infections.

**Highlights:**

- Prior maternal HIV Env vaccine trial did not assess autologous virus neutralization
- Circulating viruses isolated from mothers were tested against autologous plasma
- Maternal vaccination with HIV Env gp120/160 increased MN gp120-specific IgG binding
- Maternal HIV Env vaccine regimen did not boost autologous virus neutralization
- More potent B cell stimulation will be required to elicit autologous nAb responses

## Introduction

Despite widespread efforts to eliminate pediatric HIV infections, mother-to-child transmission (MTCT) of HIV continues to pose a significant global health challenge. With the wide availability of antiretroviral therapy (ART) for HIV-infected women during pregnancy and breastfeeding, as well as for infant prophylaxis, the rate of new HIV infections among infants have decreased by 41% from 2010 to 2018 [1]. Although approximately 82% of HIV-infected pregnant women across the globe had access to ART in 2018, there were still 160,000 newly acquired pediatric HIV infections in the same year [1]. Some of the factors contributing to these new infections are the emergence of drug-resistant HIV strains, late maternal diagnosis or presentation for prenatal care, acute infection during pregnancy or breastfeeding, and poor implementation of ART in resource-limited areas.

In the absence of ART prophylaxis during pregnancy, the MTCT transmission rate is 30-40% and can occur antepartum (*in utero*), intrapartum (during labor and delivery), or postpartum (during breastfeeding) [2]. Even with optimal implementation of antenatal triple-drug ART, breakthrough transmission can occur, with rates as high as 5% [3, 4]. In addition, recent studies have demonstrated that while ART can effectively reduce the rate of MTCT, this reduction comes at the expense of a notable increase in preterm birth and neonatal death, particularly for protease inhibitor-based regimens [4-6]. Moreover, recent reports of increased prevalence of neural tube defects in newborns associated with maternal exposure to dolutegravir-based ART at conception have raised concerns regarding toxicity of ART and highlight an urgent need for additional preventative approaches [7-10]. Thus, due to issues of ART access, adherence, incomplete efficacy, and toxicity, further strategies will be required to eliminate MTCT.

Prior studies have implicated HIV Env-specific antibody responses as being potentially protective against HIV-1 transmission. In fact, the partially effective RV144 vaccine trial of a recombinant gp120 vaccine indicated that vaccine-elicited IgG against variable loops 1 and 2 (V1V2) of gp120 was associated with decreased risk of HIV-1 heterosexual transmission [11-14]. While this particular epitope has not been implicated in protection against MTCT, maternal antibodies against both the variable loop 3 (V3) of Env gp120 and the gp41 membrane-proximal external region (MPER) have been shown to correlate with reduced risk of MTCT [15, 16]. In addition, studies have demonstrated that heterologous virus HIV-neutralizing antibodies are found more frequently or in higher titers in non-transmitting compared to transmitting mothers [17, 18]. However, there is conflicting data regarding the role of maternal antibodies in preventing vertical transmission, as other studies have failed to confirm this association between maternal Env-specific neutralizing antibodies and decreased transmission risk [2, 19]. Moreover, some studies have observed the opposite trend, reporting that transmitting women had higher concentrations of maternal IgG against the Env V3 region compared to non-transmitting women [20]. Clearly, further investigation is needed to elucidate the relationship between maternal antibody responses and risk of transmission to the infant.

We previously investigated immune correlates of vertical HIV-1 transmission in pregnant, in a large cohort of HIV clade B-infected US women from the Women and Infants Transmission Study (WITS) [21]. The results demonstrated that maternal IgG against V3, plasma neutralization of clade-matched tier 1 but not tier 2 HIV-1 variants, and the potency of the maternal plasma to block CD4 from binding to clade B HIV-1 Envs predicted reduced risk of MTCT. Interestingly, these responses were co-linear in their prediction of MTCT risk, suggesting that they may be surrogate measures for the same underlying mechanism of virus neutralization that influences infant transmission. In fact, isolated V3-specific monoclonal antibodies (mAbs) that could neutralize tier 1, but not tier 2 heterologous viruses, were able to neutralize most autologous viruses isolated from maternal plasma [21]. Moreover, it has also been demonstrated that autologous V3 and CD4 binding site (CD4bs) mAbs isolated from chronically HIV-1-infected individuals can neutralize autologous, but not heterologous, tier 2 viruses [22]. This indicates that non-broadly neutralizing antibodies can potently neutralize autologous circulating viruses, which is especially pertinent in the unique setting of MTCT, as maternal circulating viruses are the source of the vertically transmitted virus. In recent study, we also characterized vertically transmitted and non-transmitted maternal HIV Env variants in 16 mother-infant transmitting pairs from the WITS cohort, and found that, compared to maternal non-transmitted variants, the infant transmitted virus variants were significantly more neutralization-resistant to paired maternal plasma [23]. This finding suggests that autologous neutralizing antibody sensitivity may define infant transmitted/founder variants, and, therefore, boosting autologous neutralizing antibody responses in HIV infected pregnant women could be a viable immune strategy to decrease vertical transmission.

Yet, it is unknown whether vaccination of HIV-infected pregnant women with an Env vaccine would even temporarily enhance autologous virus neutralizing antibody responses. In two historic vaccine trials completed in 1993-1995 by the AIDS Vaccine Evaluation Group (AVEG) Protocols 104 and 102, safety and immunogenicity of recombinant HIV Env gp120 and gp160 (rgp120, rgp160) respectively as antigens, were tested in HIV-infected pregnant women [24]. While the Env vaccine was safe and well tolerated, there was limited enhancement of maternal immune responses against heterologous viruses in vaccinees compared to placebo recipients [24]. Importantly, because of the immunological phenomenon of original antigenic sin, it is possible that immunization with an heterologous Env vaccine may recruit memory immune cells in HIV-infected pregnant women, leading to an enhancement of their autologous virus neutralizing immune responses. In the setting of MTCT, evocation of original antigenic sin for enhancement of autologous virus neutralization could be an effective strategy to impede perinatal virus transmission. Nevertheless, whether maternal rgp120/160 vaccination enhanced neutralizing antibody responses against the maternal autologous circulating viruses remained unknown.

In this study, we sought to assess whether immunization of HIV-infected pregnant women with an alum-adjuvanted recombinant Env vaccine elicited maternal antibody responses that improved autologous virus neutralization responses. Additionally, we characterized representative maternal virus population diversity from pre- and post-immunization time points in 7 vaccinees and 3 placebo recipients and assessed the antibody binding kinetics and ability of maternal plasma to neutralize these autologous viruses. This work offers novel insights into the feasibility of enhancing maternal autologous virus neutralization and antibody responses through maternal HIV Env vaccination as an adjunctive strategy to protect the infant against HIV-1 acquisition.

## Materials & Methods

### Study Subjects

Maternal plasma samples were obtained from the AIDS Vaccine Evaluation Group (AVEG) Protocols 104 and 102, a Phase I study of safety and immunogenicity of MN rgp120 and rgp160 HIV-1 vaccines in HIV-infected pregnant women (ClinicalTrials.gov; NCT00001041). In the AVEG 104 Protocol, 26 HIV-infected pregnant women with CD4+ T cell counts >400/mm^3^ were enrolled in the second trimester of healthy pregnancy and randomized to receive either 300 µg of MN rgp120 (Genentech) with alum (n=17) as an adjuvant or alum with diluent (n=9) between 16 and 24 weeks of gestation [24]. Booster immunizations were administered monthly, until delivery, for a minimum of 3 vaccine doses and a maximum of 5 vaccine doses (Figure 1). Similarly, 2 HIV-infected pregnant women were enrolled with the same criteria in the AVEG 102 Protocol, though instead these women received either 640 µg of rgp160 (VaxSyn, MicroGeneSys) (n=1) or a placebo (n=1). Maternal plasma samples from multiple visit time points were available for 15 AVEG 104 participants (n=10 MN rgp120 vaccine, n=5 alum placebo) and 2 AVEG 102 participants (n=1 rgp160 vaccine, n=1 placebo) (Table 1).

**TABLE 1.**
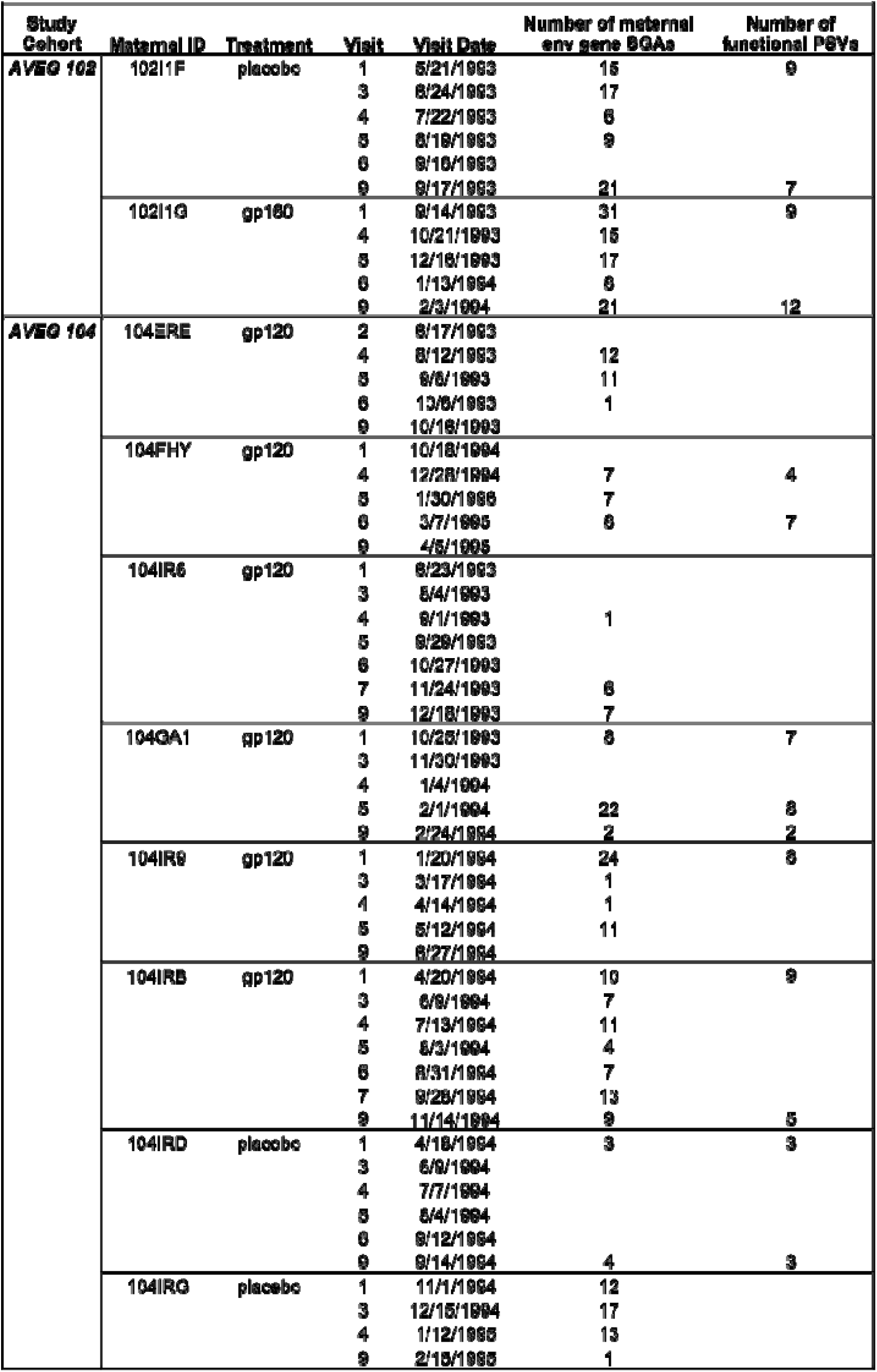
Number of maternal *env* gene sequences isolated and functional pseudoviruses produced from each maternal plasma sample.

**FIGURE 1.**
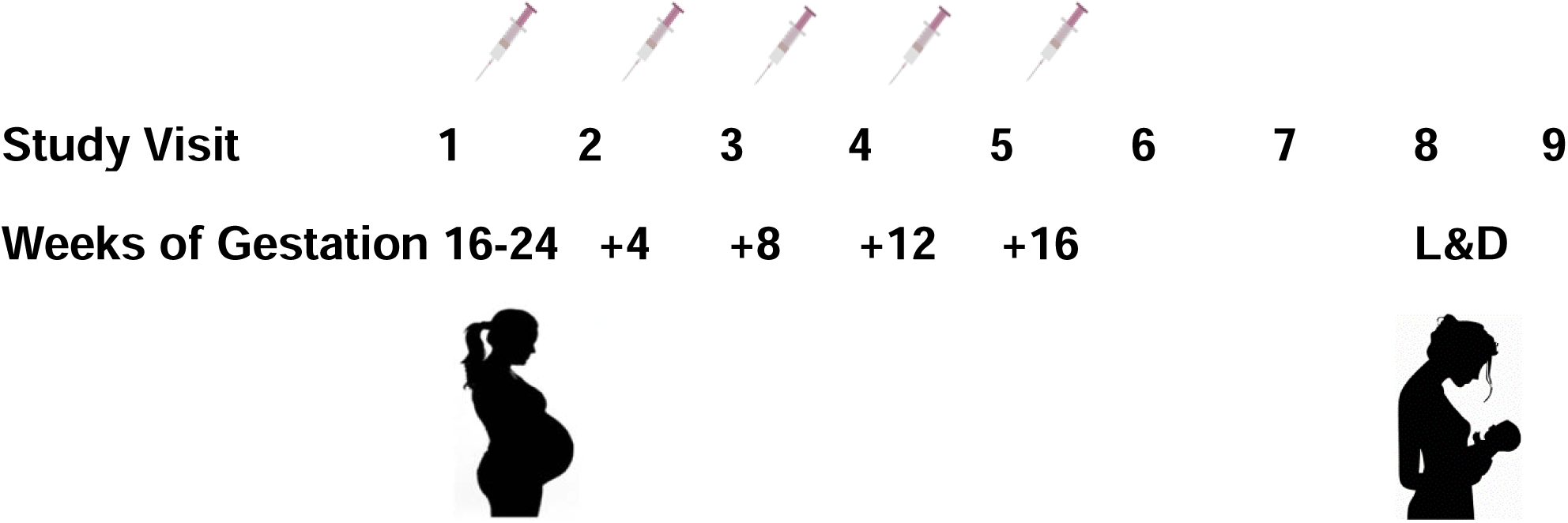
Immunization schedule in AVEG102/104 studies. Pregnant, HIV-infected women with CD4^+^ T cell counts > 400/mm^3^ were enrolled in the AVEG 102/104 studies. In the AVEG 102 protocol, women were administered 640 µg of gp160 (n=1) or placebo (n=1). In the AVEG 104 protocol, women received 300 µg of gp120 + alum (n=17) or placebo (alum + diluent) (n=9). The primary immunization was given at Visit 2, between 16-24 weeks of gestation. Monthly booster injections were subsequently given 4 weeks apart for the duration of pregnancy (Visits 3-6) for up to 5 total immunizations (median: 5; range 4-5). Visit 9 was labor and delivery.

### Ethics statement

Original study protocols AVEG 102 and 104 were approved by local institutional review boards at the seven sites involved in the original study [24]. Informed consent was obtained from all women, and also from their partners when available. In the present study, the use of de-identified maternal samples from the AVEG 102 and 104 protocol cohorts was deemed exempt by the Duke University Institutional Review Board. Moreover, in this study, individual patient identification (PTID) numbers are instead represented by PubID numbers.

### Viral RNA extraction and cDNA synthesis

Viral RNA was extracted from the plasma sample from each mother with a QIAamp Viral RNA Mini Kit (Qiagen) and subjected to reverse transcription and cDNA synthesis using 1X reaction buffer, 0.5 mM of each deoxynucleoside triphosphate (dNTP), 5 mM DTT, 2 U/mL RNaseOUT, 10 U/mL of SuperScript III reverse transcription mix (Invitrogen), and 0.25 mM antisense primer 1.R3.B3R (5’-ACTACTTGAAGCACTCAAGGCA AGCTTTATTG-3’), located in the HIV-1 *nef* open reading frame.

### Single genome amplification

Full-length envelope (*env)* genes were then amplified by nested PCR from diluted viral cDNA, as previously described [23, 25]. Briefly, cDNA was endpoint-diluted in 96-well plates (Applied Biosystems) with the goal of <30% positive amplification, in order to maximize the likelihood of obtaining a single genome. First round PCR was carried out with 1X buffer, 2 mM MgSO4, 0.2 mM of each dNTP, 0.2μM of each primer, and 0.025 U/μl Platinum Taq High Fidelity polymerase (Invitrogen) in a 20μl reaction. For the first round of PCR amplification, primer pairs were Env 5’ex (5’-TAGAGCCCTGGAAGCATCCAGGAAG-3’) and Env 3’ex (5’-TTGCTACTTGTGATTGCTCCATGT-3’), Env 5’ex and 2.R3.B6R (5’-TGAAGCACTCAAGG CAAGCTTTATTGAGGC-3’), or 07For7 (5’-AAATTAYAAAAATTCAAAATTTTCGGGTT TATTACAG-3’) and 2.R3.B6R. The following PCR conditions were used for Round 1 amplification: 1 cycle of 94°C for 2 minutes, 35 cycles of 94°C for 15 seconds, 55°C for 30 seconds, and 68°C for 3 minutes and 30 seconds for the Env 5’ex/3’ex primers (4min and 30s for Env 5’ex/2.R3.B6R, 5min and 30s for 07For7/2.R3.B6R), followed by a final cycle of 68°C for 10 minutes. Then, a second round of PCR amplification was carried out with 2μl of the first round product as template, 0.2μM of each primer, and the same PCR mixture as in round one, in a 50μl reaction. Primer pairs for second round PCR were Env 5’in (5’-TTAGGCATCTCCTATGGCAGGAAGAAG-3’) and Env 3’in (5’-GTCTCGAGATACTGCTCCCACCC-3’), Env 5’in and 2.R3.B6R (5’-TGAA GCACTCAAGG CAAGCTTTATTGAGGC-3’), or Low2c (5’-TGAGGCTTAAGCAGTGGGT TCC-3’) and VIF1 (5’-GGGTTTATTACAGGGACAGCAGAG-3’). Round 2 conditions were one cycle of 94°C for 2 minutes, 35 cycles of 94°C for 15 seconds, 55°C for 30 seconds, and 68°C for 3 minutes and 30 seconds for the Env 5’in/3’in primers (4min and 30s for Env 5’in/2.R3.B6R, 5min and 30s for Low2c/VIF1), followed by 1 cycle of 68°C for 10 minutes. Round 2 PCR amplicons were visualized using precast 1% agarose E-gels (Invitrogen), purified with the AMPure XP magnetic bead purification system (Agencourt), and sequenced for the HIV *env* gene by Sanger sequencing (Table 1).

### HIV env gene genetic analysis

Sequences were assembled using the Sequencher program (Gene Codes) and manually edited. Chromatograms were examined for sites of ambiguity, or double peaks per base read, and sequences containing multiple base peaks at a single position were marked as such and not studied further. Envelope sequences were aligned using the Gene Cutter tool available in the HIV Sequence Database of the Los Alamos National Laboratory (LANL) website (http://www.hiv.lanl.gov/content/sequence/GENE_CUTTER/cutter.html) and manually edited further in Seaview (Version 4) [26]. Phylogenetic trees were constructed using Seaview and highlighter plots were created using the Highlighter tool on the LANL website (https://www.hiv.lanl.gov/content/sequence/HIGHLIGHT/highlighter_top.html).

### Pseudovirus production and infectivity analysis

Using a previously described sequence selection algorithm [23], approximately 8-10 maternal Env variants were selected from the pre-immunization time point (Visit 1) and post-immunization time point (Visit 9) for Env pseudovirus production. Variants representing major clusters of the phylogenetic trees were selected to represent the full range of *env* genetic diversity in maternal plasma. To produce functional pseudoviruses from the HIV-1 *env* sequences, CMV promoter was added to the *env* genes by overlapping PCR as previously described [27] and the products were co-transfected with a backbone plasmid lacking the *env* gene (SG3Δenv) in 293T cells (American Tissue Culture Collection, Manassas, Virginia). 293T cells (approx. 4.5×10^6^) were seeded in a T-75 flask (Corning, Corning, NY) containing growth media (GM) (Dulbecco’s modified Eagle’s medium (DMEM)-10% fetal bovine serum (FBS)-1% penicillin-streptomycin containing HEPES, ThermoFisher, Waltham, MA) and incubated overnight at 37°C and 8% CO_2_. 4μg of Env DNA containing CMV promoter was combined with 4μg of SG3Δenv backbone and FuGene 6 transfection reagent (Roche Diagnostics) was added as per manufacturer’s instructions. The mixture was then added to the T-75 flask and it was incubated at 37°C for 48 hours. Supernatant containing pseudovirus was harvested and stored at −80°C with a final concentration of 20% FBS. To measure the infectivity of the pseudoviruses, 20μl of pseudovirus was added in duplicate to a 96-well flat bottom plate and then 100μl TZM-bl cells (catalog no. 8129; NIH AIDS Reagent Program; from John Kappes and Xiaoyun Wu) were added (10,000 cells/100μl GM with 10μg/ml of DEAE-dextran). After a 48-hour incubation at 37°C and 8% CO_2_, 100μl of culture medium was removed and 100μl of Bright-Glo luciferase reagent (Promega) was added. The mixture was incubated for 2 minutes at 25°C, 100μl was subsequently transferred to a 96-well black plate, and luminescence was measured immediately on a Victor X3 multilabel plate reader (PerkinElmer).

### TZM-bl neutralization assay

Neutralization of autologous pseudoviruses by maternal plasma was measured using a luciferase (Luc) reporter gene assay in TZM-bl cells (catalog no. 8129; NIH AIDS Reagent Program; from John Kappes and Xiaoyun Wu), as previously described [28]. Before performing the assay, plasma was heat inactivated by incubating for 30 minutes at 56°C. Plasma samples were added at a starting dilution of 1:20 and diluted threefold serially. Then, the plasma samples were incubated with virus for 1 hour at 37°C. TZM-bl cells were added, and the mixture was incubated for 48 hours. Luminesence was then measured using the Bright-Glo luciferase reagent and Victor X3 luminometer and luminescence values used to calculate the ID_50_, or dilution at which relative luminescence units (RLU) were reduced by 50% compared to virus control wells. VRC01 was used as a positive control in each experiment and murine leukemia virus (SVA.MLV) served as a negative control for the assay [29].

### Binding antibody multiplex assay (BAMA)

HIV-1 Env-specific IgG responses in maternal plasma against a panel of HIV-1 antigens were detected using a customized BAMA, as previously described [30]. HIV-1 antigens were covalently coupled to carboxylated fluorescent beads (Bio-Rad Laboratories) and IgG binding to the bead-coupled antigens was measured. The antigen panel for IgG BAMA assays included biotinylated linear V3 loop peptide V3.B (Bio V3 B) and the following 4 proteins: MNgp120, Gp70 B.MN V3, Gp70 B.CaseA_V1V2, and HIV-1 MN recombinant gp41 (REC MN gp41, ImmunoDiagnostics) (Table S1). The antigen-coupled beads were incubated with diluted plasma samples (1:100 for MNgp120, Gp70 B.MN V3, Gp70 B.CaseA_V1V2; 1:2000 for V3.B and REC MN gp41) for 30 minutes at room temperature (20-25°C). HIV Env-specific IgG was then detected with phycoerythrin (PE)-conjugated mouse anti-human IgG (Southern Biotech, Birmingham, AL) at 2 μg/ml. Beads were washed, resuspended, and acquired on a Bio-Plex 200 instrument (Bio-Rad Laboratories). Blank beads were used to account for non-specific binding and HIV immunoglobulin (HIVIG) was used as a positive control for all assays. The magnitude of antibody binding to the panel of HIV-1 Env antigens was measured as mean fluorescent intensity (MFI). MFI values for conformational antigens containing gp70 were background-adjusted by subtracting the MFI values of gp70 MulV. All MFI values were background-adjusted by subtracting the MFI values of coupled beads without sample. A positive HIV Env-specific antibody response was considered to be an MFI > 100. The criteria for reporting sample MFI values included coefficient of variation ≤20% with a bead count ≥100 for each sample. All assays tracked the 50% effective concentration and maximum MFI of the positive control HIVIG and protein standards CH58, B12, and 7B2 by Levey-Jennings charts to ensure data consistency.

### Statistics

We tested for differences in antibody binding and neutralization responses between placebo and vaccinees using 2-sided Wilcoxon tests comparing the values at the first visit to those at the last visit. Because time intervals between visits differed across study subjects, we also tested the change per day between visits. All statistical analyses and graphs were produced using R [31].

## Results

### Env-specific antibody binding responses in vaccinees compared to placebo recipients

The magnitude of HIV Env epitope-specific IgG responses prior to and following Env vaccination in HIV-1 infected vaccinated women and placebo recipients was assessed by BAMA. We measured maternal vaccine-elicited responses against clade B MN gp120 protein (Env matched to the vaccine immunogen), gp70 V1V2 protein, and a linear V3 peptide. When comparing the change in clade B MN gp120-specific binding responses between the first and last visit among study participants, all placebo recipients (n=6) did not show increase in binding, whereas 8 out of 11 vaccinees showed increase in binding over time. Statistically, the overall increase in binding to MN gp120 was significantly higher in vaccinees compared to placebo recipients (p=0.027 by Wilcoxon test, Figure 2). To account for differences in the timing of visits for each mother, we calculated the change in MN gp120-specific binding response per day between the first (Visit 1, 4) and last visit (Visit 9). Per day increase in clade B MN gp120-specific binding responses between the first and last visit was statistically significantly higher in vaccinees as compared to placebo recipients (p=0.015 by Wilcoxon test, Figure S1). We observed more than 3-fold increase in antibody binding responses against MN gp120 in 1/11, 2/11, and 3/11 vaccinees against antigens MN gp120, linear V3.B, and gp70 V1V2, respectively. No change in change in antibody binding responses was observed in placebo recipients except in one against gp70 V1V2 (Figure 3, Figure S2).

**FIGURE 2.**
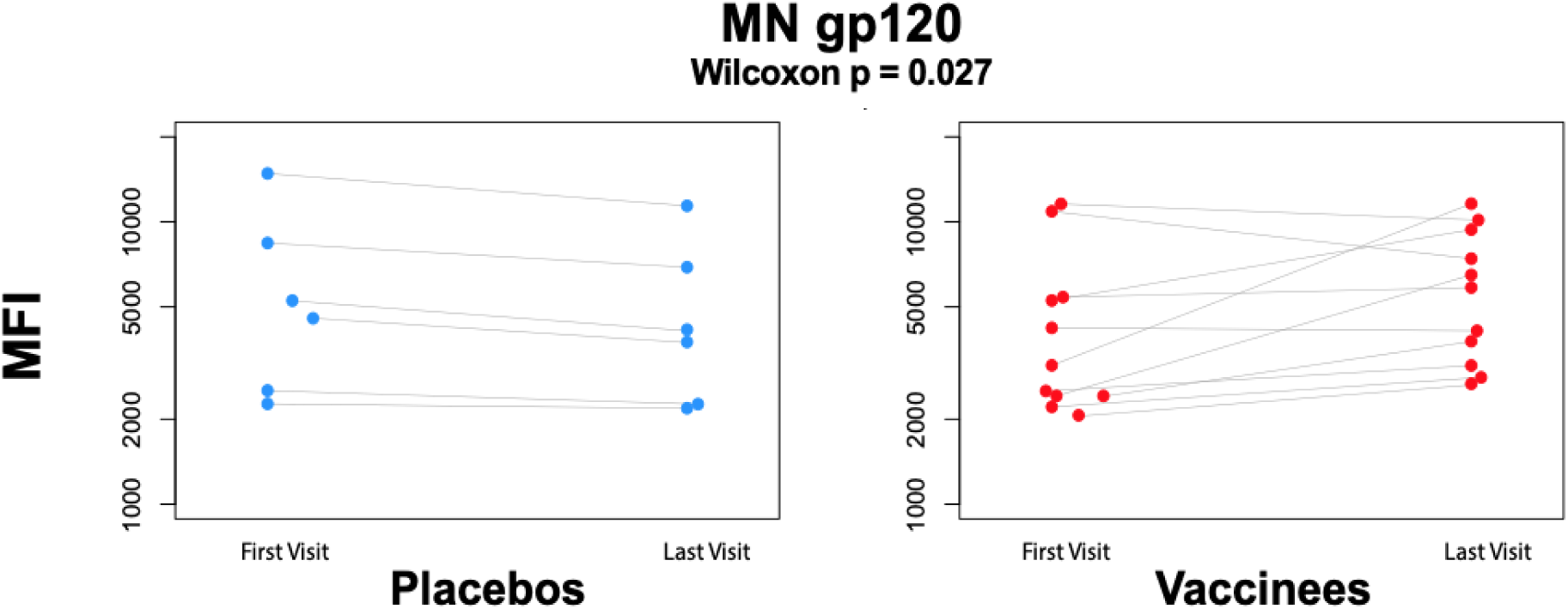
MN gp120-specific binding in vaccinees and placebo recipients at first and last visit. Comparison of changes in gp120-specific binding from the first to the last visit between vaccinees (red) and placebo (blue). The between-visits change in gp120-specific binding was statistically significantly higher in vaccinees (p=0.027 by 2-sided Wilcoxon test). Light gray lines link the same mother.

**FIGURE 3.**
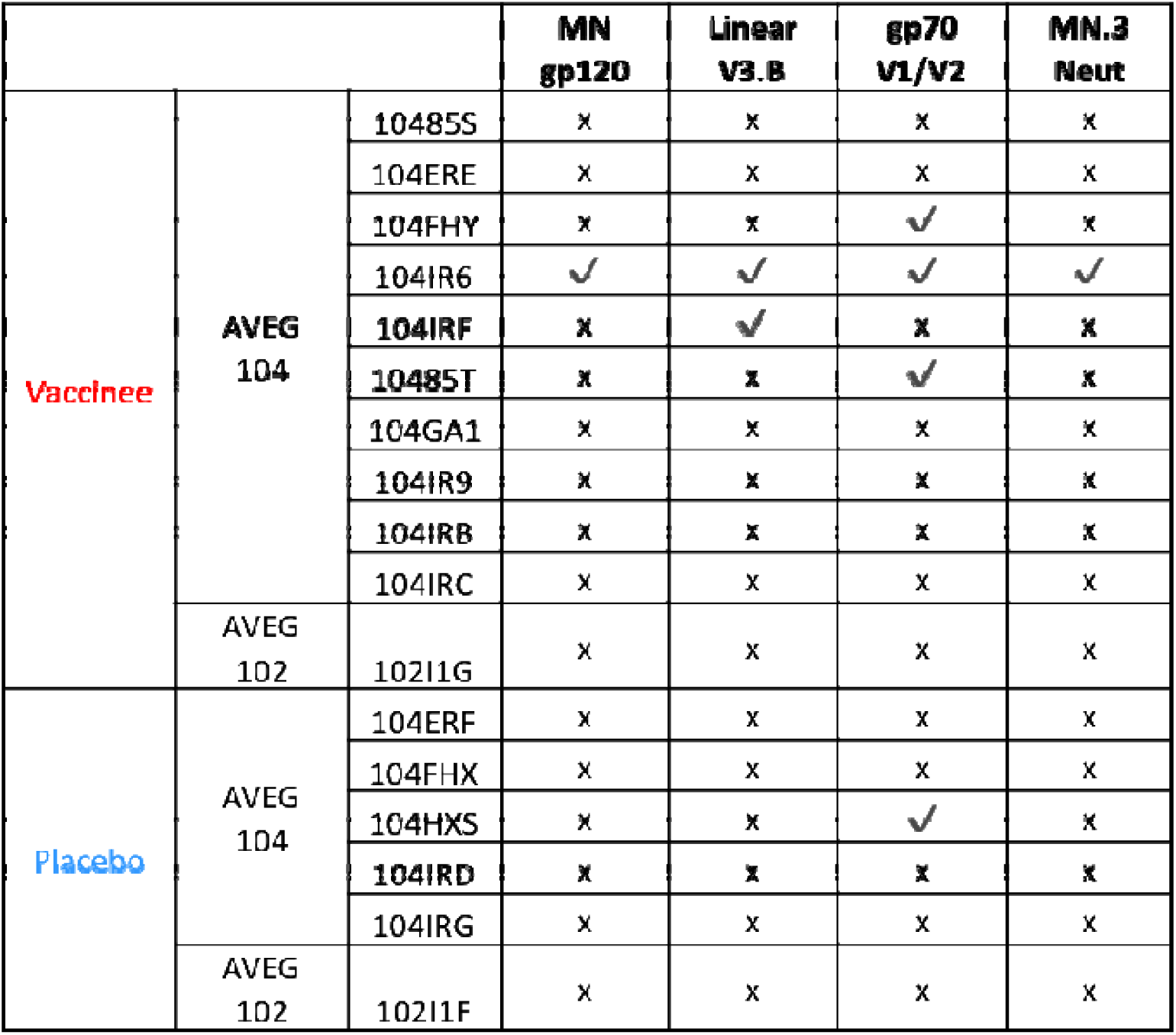
Change in antibody binding response against MN.3 gp120, linear V3.B, gp70 V1V2 and neutralization response against MN.3 before and after vaccination. Vaccinees (red) or placebo recipients (blue). Check marks indicate a three-fold increase.

### Autologous virus neutralizing antibody responses in vaccinees compared to placebo recipients

To determine if gp120 or gp160 vaccination enhanced functional, virus-specific neutralizing antibody responses, we assessed the ability of maternal plasma to neutralize autologous virus variants isolated from plasma collected in early pregnancy before vaccination and late pregnancy after vaccine boosting. There was no difference between vaccinees and placebo recipients in the ability of maternal plasma collected at delivery to neutralize autologous virus populations isolated from early and late pregnancy visits (Figure 4A-B, Figure S3). Moreover, the difference in the ability of maternal plasma collected at the first visit (Visit 1,4) versus last visit (Visit 9) to neutralize early pregnancy plasma viruses (Visit 1, 4) was comparable between vaccinees and placebo recipients (Figure 4C-D).

**FIGURE 4.**
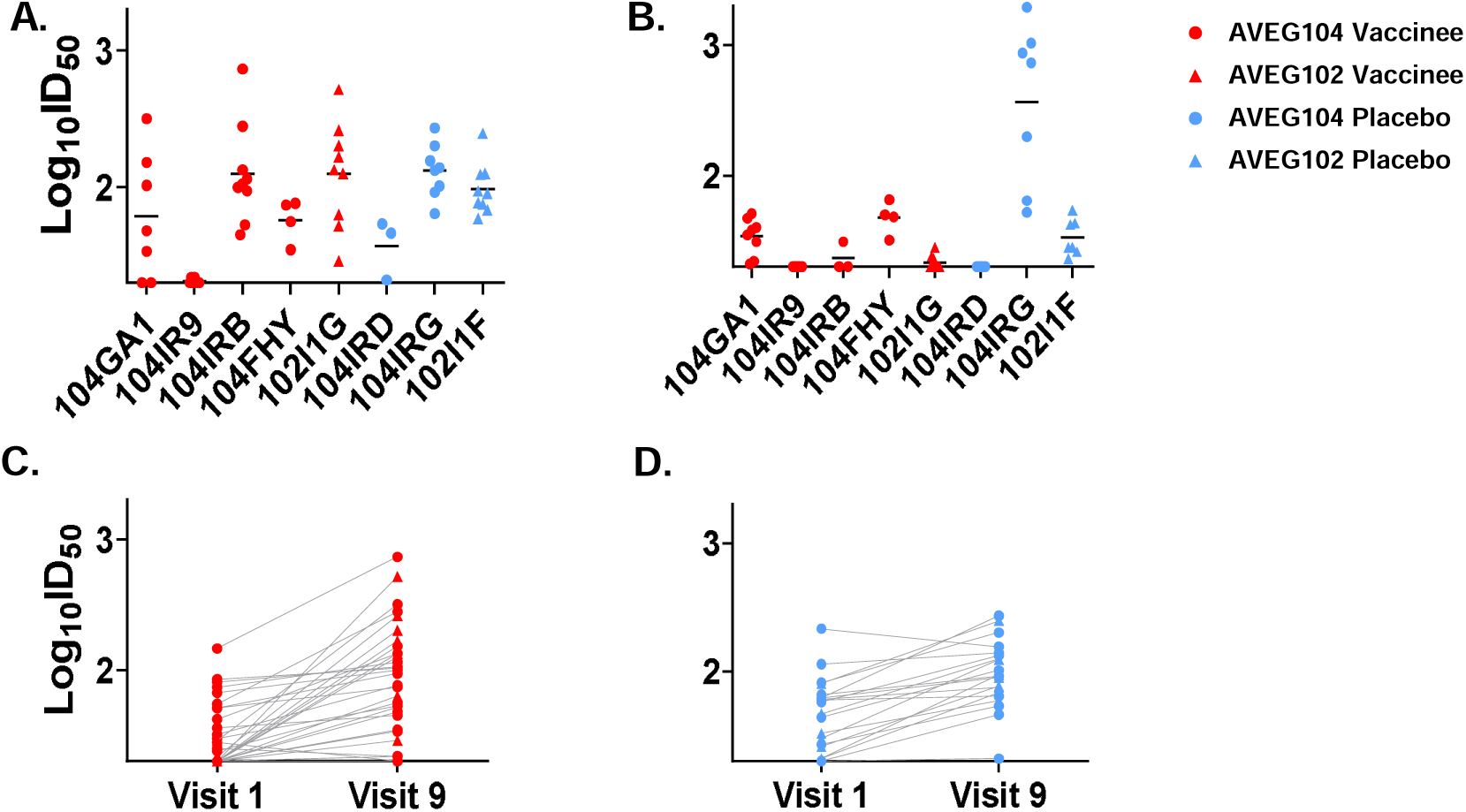
Neutralization of viruses isolated from vaccine and placebo recipient plasma during early and late pregnancy by autologous maternal plasma collected at delivery. For each vaccine and placebo recipient, the neutralization potency of maternal plasma at delivery was assessed against the early pregnancy (Visits 1,4) (A) and late pregnancy (Visits 5-9) (B) autologous virus populations. The left y-axis depicts neutralization potency, in log_10_ID_50_. Study participants are displayed on the X axis (A,B). AVEG104 study participants are depicted with circles; AVEG 102 study participants with triangles. Vaccine recipients are shown in red; placebos in blue. Bars represent geometric means. Change in neutralization potency of autologous maternal plasma from Visits 1 (pre-immunization) and 9 (delivery) are shown against early pregnancy plasma viruses (C,D), with the exception off autologous viruses isolated from Visit 4 in mother 104FHY.

Additionally, we tested the ability of maternal plasma collected from the pre-vaccination visit (Visit 1), the booster visits (Visits 3, 4, 5, & 6), and the post-vaccination visits (Visits 7 & 9) to neutralize autologous virus populations from the pre-vaccination or early pregnancy visit (Visit 1, 4) (Figure 5). Interestingly, there appears to be a boost over time in neutralization potency against autologous virus populations isolated from the pre-vaccination visit in the gp160 vaccinee as compared to the corresponding placebo recipient, yet this trend was only observed in the one vaccinee from which samples were available for testing. Taken together, there was no significant change in autologous virus neutralization potency over time for Env vaccinees as compared to placebo recipients.

**FIGURE 5.**
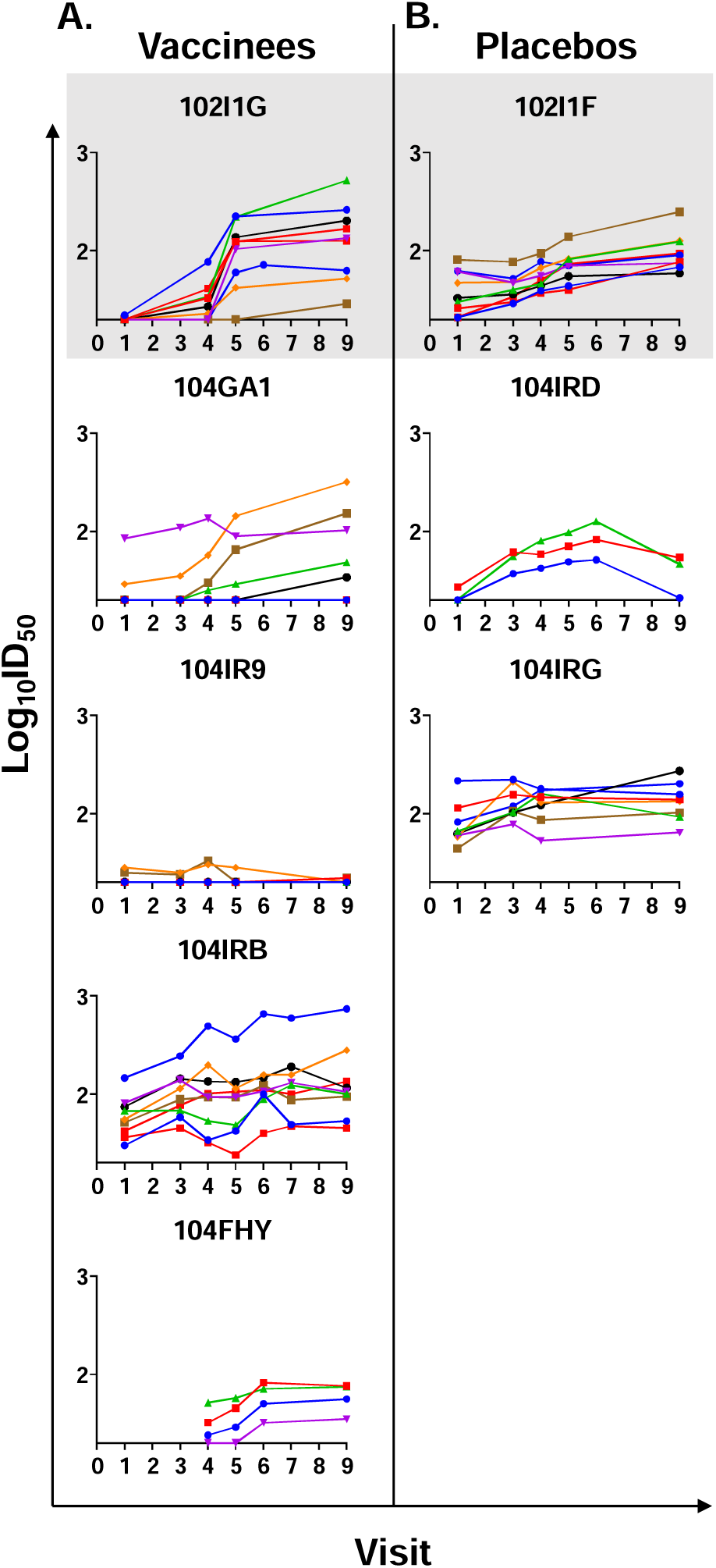
Neutralization potency of autologous maternal plasma against plasma viruses isolated from early pregnancy. Maternal plasma from pre-immunization (Visit 1), booster visits (Visits 3, 4, 5, & 6) and post-immunization (Visits 7 & 9) was tested against individual virus variants from Visit 1 (except for 104FHY; virus variants are from Visit 4). Each colored line represents a different virus. The left y-axis depicts neutralization potency, in log_10_ID_50_. AVEG 102 study participants are shown on the top row, shaded in gray. With additional gp160 doses, there is a greater boost in neutralization potency against autologous viruses in the gp160 vaccine recipient compared to the placebo. There does not appear to be an increase in autologous virus neutralization with gp120 vaccination compared to placebo.

### Plasma HIV env gene sequence diversity in vaccinees compared to placebo recipients

Through single genome amplification, we obtained 282 total HIV *env* gene sequences from vaccinees (n=7) and 118 total HIV *env* gene sequences from placebo recipients (n=3) (Table 1). To characterize viral evolution between visits in study participants, we measured viral diversity through the mean pairwise Hamming distance, which was calculated as the number of mutations between all pairs of *env* sequences isolated from one sample divided by the total number of bases in the alignment. The changes in viral diversity between visits observed in the placebo recipients were not significantly different from those observed in the vaccinees (Figure 6). Yet, in a per day analysis of mean Hamming distance since the pre-vaccine visit, we observed a trend in which vaccinees consistently demonstrated lower HIV *env* gene sequence diversity than placebo recipients (Figure S4). However, this trend did not reach significance, potentially due to limited statistical power given the small sample size.

**FIGURE 6.**
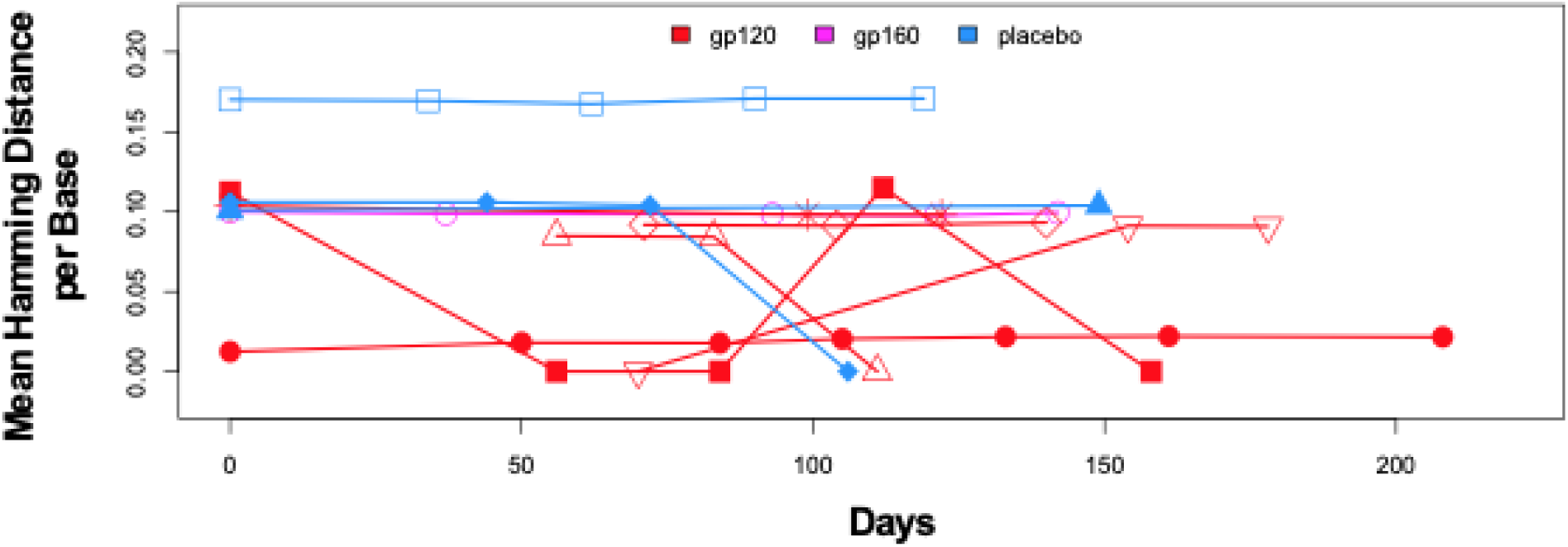
Comparison of change in intersequence Hamming distance per base pair of *env* sequences obtained from vaccinees (n=7) and placebo recipients (n=3) across study visits in days. Each shape is an individual mother. Gp120 (red), gp160 (pink), placebo (blue).

## Discussion

Unlike other modes of HIV transmission, MTCT is a unique setting since it occurs in the presence of pre-existing humoral immunity. Maternal IgG antibodies that are passively acquired by the fetus through transplacental transfer during pregnancy and by the infant through breastfeeding are known to provide critical protection against perinatal pathogens for neonates during the first few months of life [32]. Despite the potential source of transmitted virus in a mother-infant pair being confined to the maternal autologous circulating virus pool, our limited understanding of how maternal antibody responses that coevolved with circulating virus populations impact vertical HIV transmission remains a major barrier in developing a maternal immunization strategy for the prevention of MTCT. Yet, there are reasons to believe that enhancing the ability of maternal plasma to neutralize co-circulating viruses may impede vertical virus transmission. Thus, augmentation of maternal autologous virus neutralizing antibody responses through vaccination during pregnancy is a potentially viable strategy to further reduce the rate of MTCT in synergy with ART.

In this study, we investigated the induction of neutralizing antibodies against autologous circulating viruses in the AVEG 102 and 104 maternal HIV Env vaccine trials (1993-1995), which assessed the safety and immunogenicity of alum-adjuvanted gp120 and gp160 subunit vaccines in HIV-infected, pregnant women. Importantly, while the Env vaccines were reported to be safe and immunogenic, the ability of the vaccines to raise neutralizing responses against maternal circulating autologous viruses was not assessed [24]. With the development of novel techniques to isolate and analyze single HIV variants from plasma, as well as more sensitive, bead-based binding antibody detection, we are now better equipped to answer this important question. Another limitation to the present study is incomplete information from the original study related to which subset of participants may have received antiretroviral therapy, as recommendations for clinical management of HIV-infected pregnant women were updated during the study following the first recognition that zidovudine was effective in decreasing the risk of MTCT of HIV [33]. Additionally, sample quality and restricted availability of time points limited the successful isolation of sufficient number of single genome amplicons from each participant for larger analyses. While the small sample size of 11 vaccinees and 6 placebo is a caveat to this study, these cohorts of HIV-infected pregnant women enrolled in a Phase I HIV Env subunit vaccine clinical trial represented a unique opportunity to understand the ability of HIV Env vaccination to enhance neutralizing antibody responses in the setting of MTCT.

Although enhanced autologous virus neutralization was not observed in the AVEG 102 and 104 study cohorts, our work offers key insight in that it suggests more potent B cell stimulation must be achieved for an HIV Env vaccination to be effective in boosting autologous neutralizing antibody responses in pregnancy. There are valuable lessons to be gained and key opportunities for improvement in trial design based on our analysis of this first Phase I trial of HIV Env subunit recombinant gp120 and gp160 protein vaccines adjuvanted with alum in HIV-infected, pregnant women. While the first formal recognition that zidovudine (ZDV) was effective in reducing the risk of MTCT occurred during the AVEG 102 and 104 trial periods [33], next-generation maternal HIV Env vaccination strategies should be developed within the contemporary context of widespread availability of ART for pregnant women. Consequently, studies or trials of future maternal vaccine regimens aimed at preventing vertical transmission of HIV should model the conditions of antiretroviral therapy and viral suppression during pregnancy.

Though the majority of vaccines currently licensed for human use in the United States are formulated with an aluminum-based adjuvant, alternative adjuvant selection may play a critical role in the elicitation of protective humoral responses against HIV transmission [34, 35]. In a systematic comparison of the ability of adjuvant formulations, including Toll-like receptor (TLR) agonists, to induce antibody binding, neutralizing, and ADCC responses against transmitted/founder HIV-1 envelope gp140 (B.63521) in rhesus macaques, Moody et al. demonstrated that combination of a TLR7/8 agonist with a TLR9 agonist in a squalene-based oil-in-water emulsion resulted in enhanced HIV Env-specific antibody responses [36]. These potent TLR-based adjuvants may be exploited to synergize with highly antigenic HIV immunogens to induce more potent and durable protective antibody titers. During the 2009 influenza A (H1N1) pandemic, an H1N1 vaccine in combination with a novel squalene-based AS03 adjuvant was safely used among pregnant women in Norway and resulted in reduced risk of both influenza diagnosis and fetal death [37]. Thus, the demonstrated safety profile of AS03 adjuvant among pregnant women has opened the door for implementation of novel adjuvants beyond alum in pregnancy.

Moreover, the nature of vertical transmission of HIV necessitates a vaccination strategy that can mount a protective immune response specific to the autologous circulating virus pool of each HIV-infected mother. Thus, the choice of vaccine immunogen must reflect the broad diversity of HIV strains circulating today, specifically clade C and B virus subtypes prevalent in Sub-Saharan Africa and the United States and Europe, respectively. One potential approach to overcome this barrier is to employ a multi-clade (B/C) HIV Env immunogen that would leverage the immunological phenomenon of original antigenic sin to provoke an anamnestic response against maternal autologous circulating viruses.

Additionally, future studies may explore the potentially protective role of antibody-mediated effector functions beyond neutralization in reducing the risk of MTCT of HIV. Notably, a previous study by Overbaugh et al. suggested that HIV Env-specific IgG-mediated antibody dependent cell cytotoxicity (ADCC) activity in breastmilk correlates with reduced risk of postnatal vertical transmission [38]. Moreover, we have previously demonstrated that passive infusion with a cocktail of non-neutralizing antibodies provided partial protection against postnatal SHIV acquisition in an infant non-human primate oral challenge model [39]. Further investigation into the potentially protective roles of other Fc-mediated effector functions including antibody dependent cell phagocytosis (ADCP) and complement activation activity in the setting of MTCT of HIV is warranted. Perhaps elicitation of the full breadth of polyfunctional antiviral activity of the humoral immune response, not only autologous neutralization response, will be the critical target of maternal HIV vaccine design to eliminate MTCT.

## Conclusion

In this study, we assessed the autologous virus neutralization responses of maternal plasma collected at delivery against circulating viruses isolated from early and late pregnancy in HIV-infected women vaccinated with an HIV Env subunit recombinant gp120/160 adjuvanted with alum from historical AVEG 102 and 104 Phase I trials. We found that vaccination of HIV-infected pregnant women with recombinant MN gp120 or gp160 adjuvanted with alum boosted HIV Env-specific antibody binding responses between the first and last visit against clade B MN.3 gp120, the original vaccine antigen, compared to placebo recipients. However, vaccination failed to augment the ability of maternal plasma collected at delivery to neutralize clade B MN.3 virus, a tier 1 heterologous virus, between the first and last visit. Maternal HIV Env vaccination did not enhance the ability of maternal plasma collected at delivery to neutralize autologous viruses isolated from early pregnancy. Moreover, vaccination had no evident impact on viral diversity. These findings indicate that further optimization in choice of vaccine immunogen and adjuvant will be necessary to effectively augment autologous virus neutralization responses in HIV-infected pregnant women to synergize with ART and reduce MTCT of HIV.

## Abbreviations

(nAb): Neutralizing antibody
(Env): envelope protein
(AVEG): AIDS Vaccine Evaluation Group
(SGA): single genome amplification
(*env*): envelope gene
(MTCT): mother-to-child transmission
(ART): antiretroviral therapy
(V1V2): variable loops 1 and 2
(V3): variable loop 3
(MPER): membrane-proximal external region
(WITS): Women and Infants Transmission Study
(mAbs): monoclonal antibodies
(CD4bs): CD4 binding site
(rgp120): recombinant HIV Env gp120
(rgp160): recombinant HIV
(BAMA): binding antibody multiplex assay
(HIVIG): HIV immunoglobulin
(MFI): mean fluorescent intensity
(ZDV): zidovudine
(TLR): Toll-like receptor
(ADCC): antibody dependent cell cytotoxicity

## Author Contributions

**Eliza D. Hompe**: Conceptualization, Data curation, Formal analysis, Funding acquisition, Investigation, Methodology, Writing – original draft, Writing – review & editing. **Jesse F. Mangold**: Data curation, Formal analysis, Visualization, Writing – original draft. **Joshua A. Eudailey**: Data curation, Formal analysis, Investigation, Writing – review & editing. **Elena E. Giorgi**: Formal analysis, Methodology, Software, Writing – review & editing. **Amit Kumar**: Investigation, Methodology, Writing – review & editing. **Erin McGuire**: Investigation. **Barton F. Haynes**: Writing – review & editing. **M. Anthony Moody**: Writing – review & editing. **Peter F. Wright**: Resources, Writing – review & editing. **Genevieve G. Fouda**: Resources, Writing – review & editing, Supervision. **Feng Gao**: Resources, Writing – review & editing, Supervision. **Sallie R. Permar**: Conceptualization, Methodology, Resources, Writing – review & editing, Supervision, Project administration, Funding acquisition. All authors gave final approval of the manuscript to be submitted. All authors attest they meet the ICMJE criteria for authorship.

## Acknowledgements

We thank the women who participated in the AIDS Vaccine Evaluation Group 102 and 104 trials and the HIV Vaccine Trials Network for supporting identification and handling of the maternal plasma specimens to conduct the present study. We would also like to acknowledge Hanna Itell, Grace Li, and Maria Dennis for their assistance with BAMA assays.

---

**FIGURE S1.**
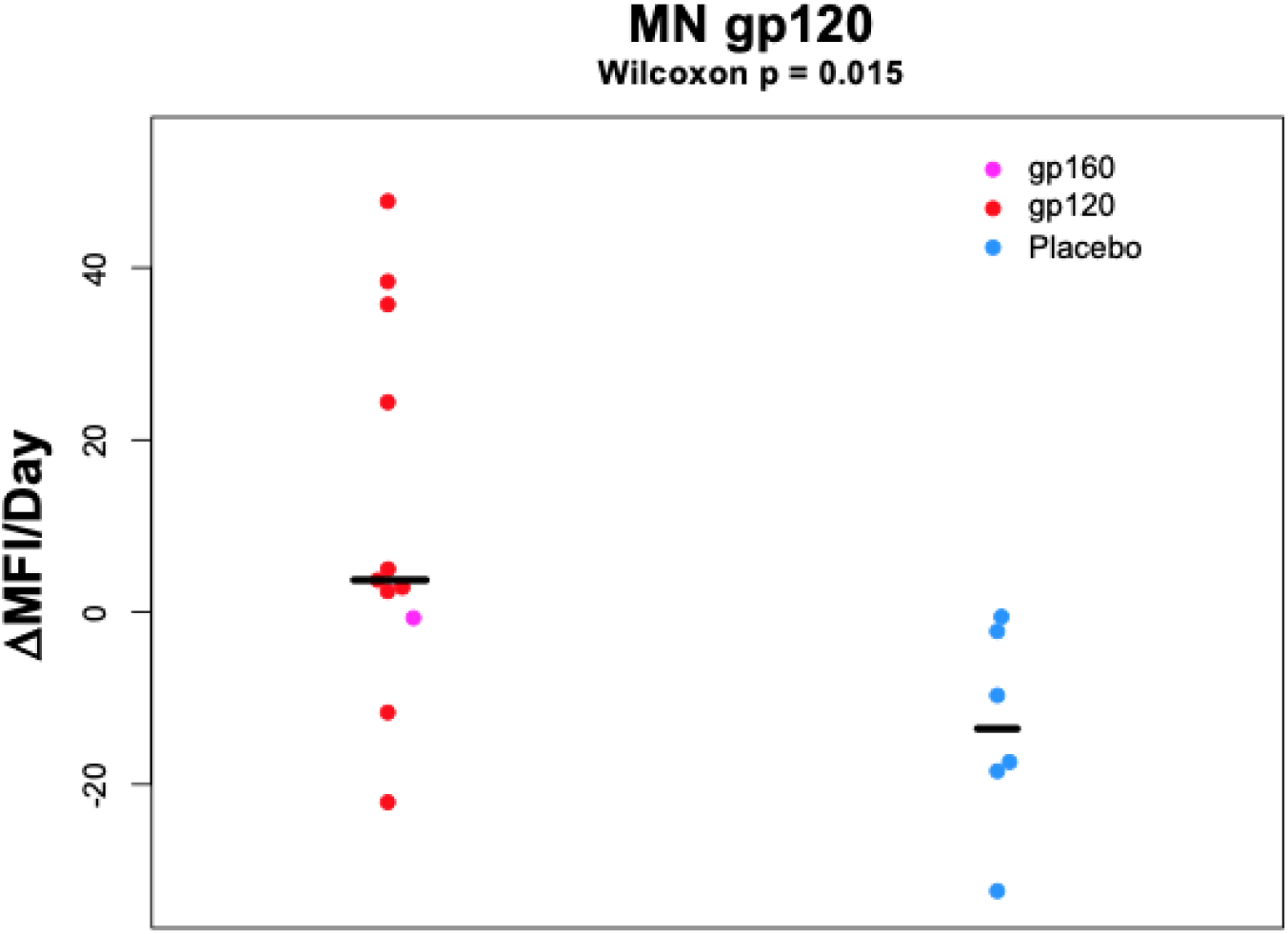
Change in MN gp120-specific binding per day in vaccinees and placebo recipients between first and last visit. Comparison of changes in gp120-specific binding per day from the first to the last visit between vaccinees (red or pink) and placebo (blue). The gp120 specific binding increase per day was higher in vaccinees compared to placebo (p=0.015 by 2-sided Wilcoxon test).

**FIGURE S2.**
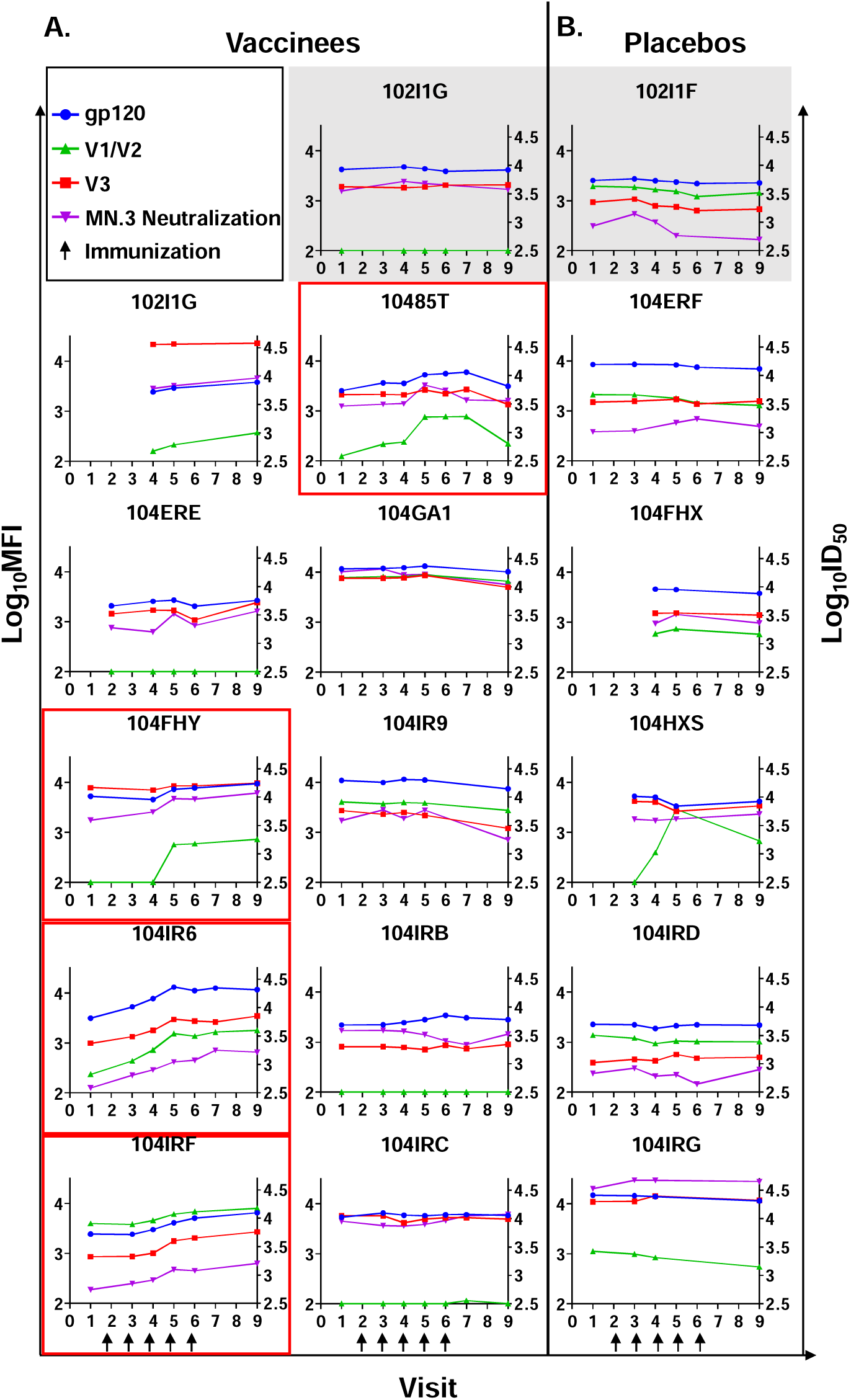
HIV Env gp120, V1V2, and V3-specific IgG binding and neutralizing Ab responses in HIV-infected, pregnant women following immunizations with MN gp120 or gp160. The left y-axis depicts binding antibody response, in log_10_MFI, and gp120 binding responses are indicated by the blue lines, V3 responses by the red lines, and V1V2 responses by the green lines. The right y-axis depicts neutralization potency, in log_10_ID_50_, and is indicated by the purple lines. AVEG 102 study participants are shown on the top row, shaded in gray. Red boxing indicates 3-fold increase in gp120, V1V2, V3, or neutralizing antibody responses over time.

**FIGURE S3.**
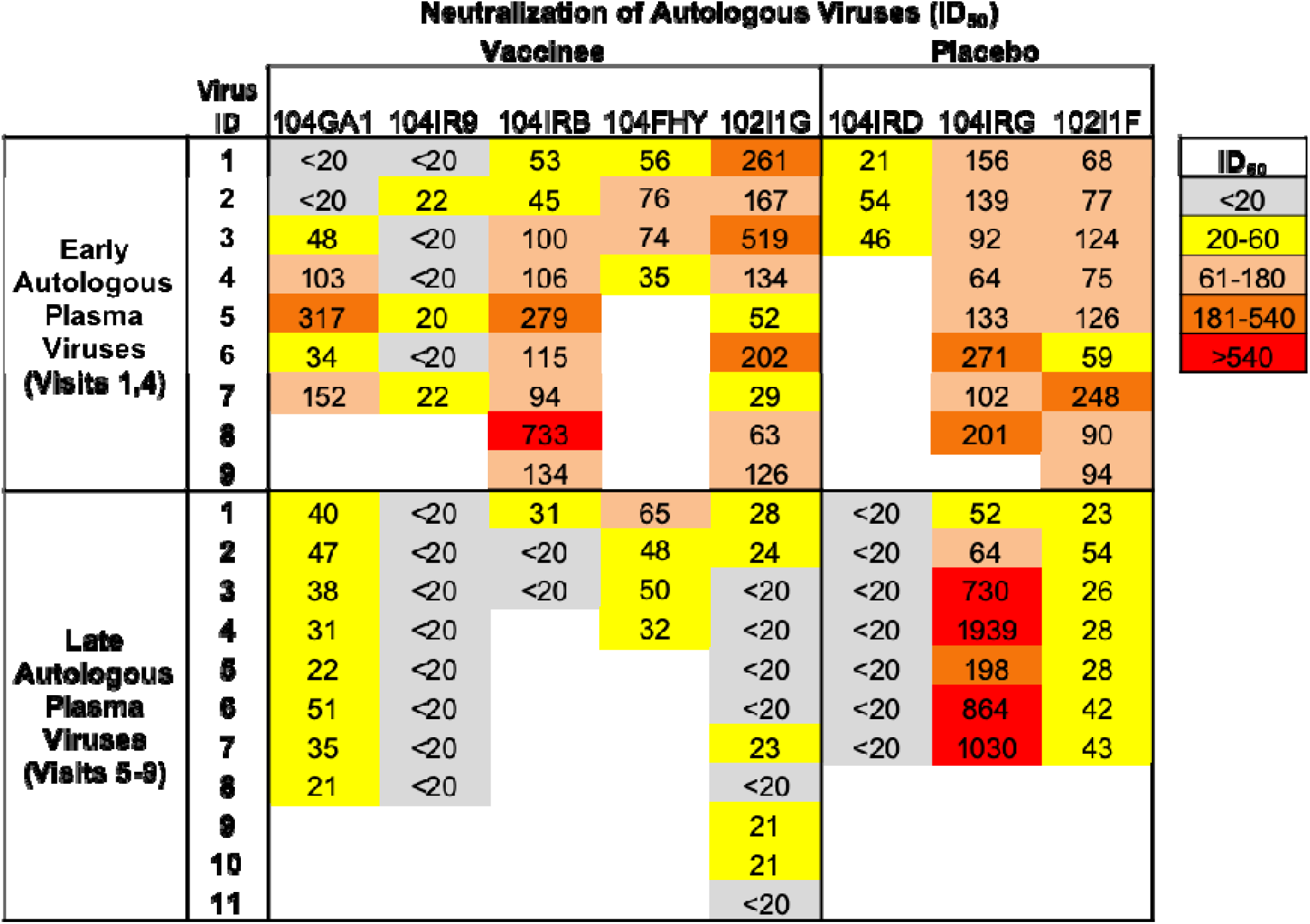
Neutralization of viruses isolated from vaccine and placebo recipient plasma during early and late pregnancy by autologous maternal plasma collected at delivery. For each vaccine and placebo recipient, the neutralization potency of maternal plasma at delivery was assessed against the early (Visits 1,4) and late pregnancy (Visits 5-9) autologous virus populations. Higher ID_50_ (darker color) represents greater neutralization potency.

**FIGURE S4.**
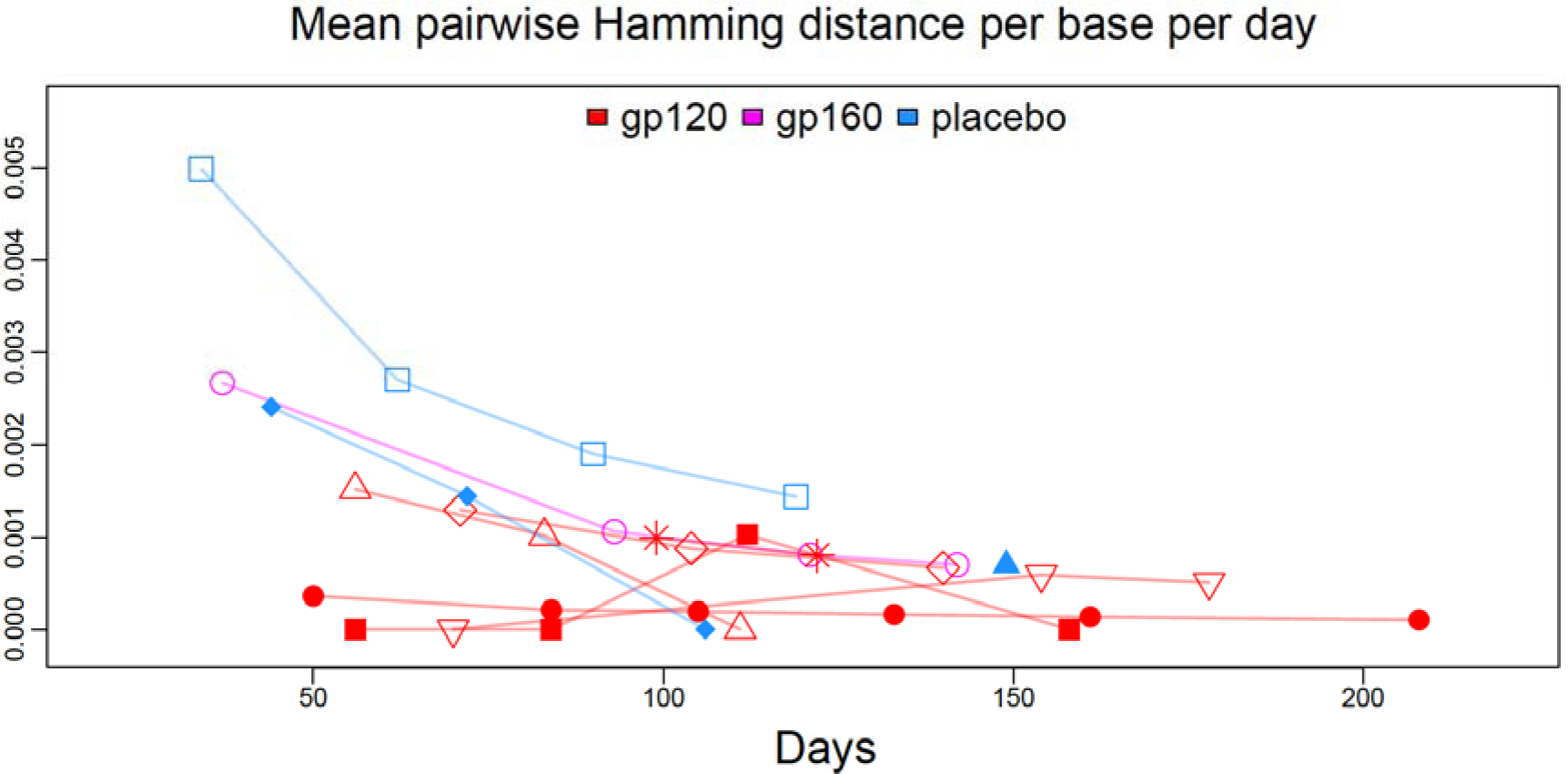
Comparison of change in intersequence Hamming distance per base pair per day of *env* sequences obtained from vaccinees (n=7) and placebo recipients (n=3) across study visits. Each shape is an individual mother. Gp120 (red), gp160 (pink), placebo (blue).

**TABLE S1.**
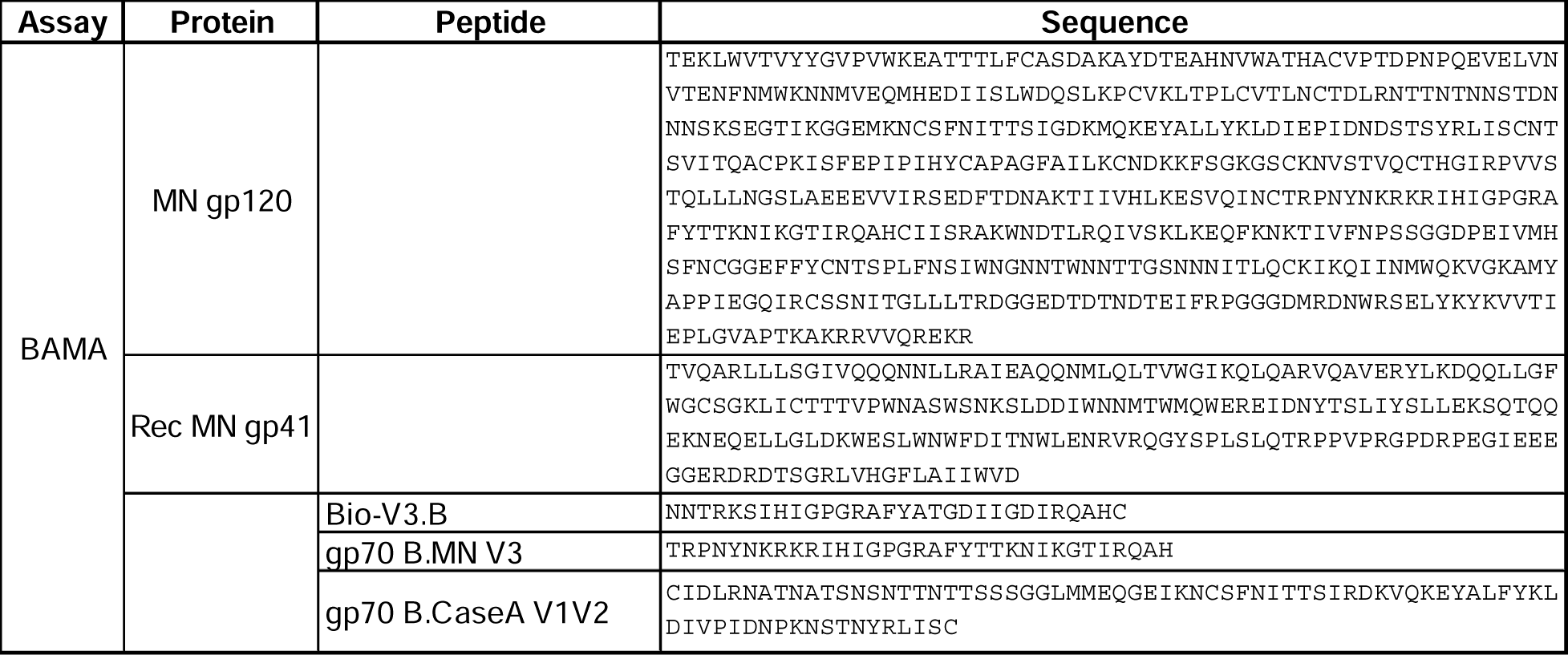
Amino acid sequences for antigens used in AVEG102/104 plasma binding antibody multiplex assays (BAMA).

